# Characterization of the bacterial microbiome associated with centrohelid heliozoans from aquatic environments using full-length 16S rRNA PacBio sequencing

**DOI:** 10.64898/2026.03.19.712920

**Authors:** Elena A. Gerasimova, Alexander S. Balkin, German A. Sozonov, Tatyana A. Chagan, Elizaveta I. Kaleeva, Ruslan Kasseinov, Darya V. Poshvina

## Abstract

Centrohelid heliozoans are a monophyletic group of free-living, ubiquitous, predatory protists widely distributed in aquatic and soil ecosystems. Centrohelids are known as cytotrophic protists that feed on bacteria, algae, and small unicellular eukaryotes. While algal and chloroplast symbioses have been documented in this group, their bacterial associations remain largely unexplored. In this study, we characterize the bacterial communities associated with centrohelids isolated from freshwater habitats using full-length 16S rRNA PacBio sequencing. Amplicon sequencing revealed 5 phyla, 6 classes, and 58 genera in the bacterial communities associated with seven centrohelid isolates. Alphaproteobacteria, Bacteroidia, and Gammaproteobacteria were the most abundant classes, while *Arcicella*, *Sphingobium*, *Pseudomonas*, *Sphingomonas*, *Azospirillum*, *Shinella*, *Flavobacterium*, *Variovorax*, and *Rhodococcus* were the most abundant genera. Notably, *Arcicella*, *Variovorax*, *Sphingobium*, and *Pseudomonas* constituted the core microbiome. Unexpectedly, we detected bacteria known as opportunistic pathogens, providing the first evidence that centrohelids may serve as environmental reservoirs for bacteria with pathogenic potential (e.g., *Acidovorax*, *Acinetobacter*, *Anaerococcus*, *Bosea*, *Corynebacterium*, *Escherichia*, *Moraxella*, *Mycobacterium*, *Prevotella*, *Pseudomonas*, *Ralstonia*, and *Sphingomonas*). In addition, this study provides the first evidence of Rickettsiaceae associations with centrohelids.

**IMPORTANCE:** This study reveals that centrohelid heliozoans, ubiquitous microbial predators, harbor diverse and host-specific bacterial communities. Critically, we show they can serve as environmental reservoirs for bacteria with pathogenic potential, a role previously overlooked outside of model protist groups. These findings expand our understanding of pathogen ecology, suggesting that a wider range of protists may contribute to the persistence and dispersal of opportunistic pathogens in aquatic ecosystems.

## INTRODUCTION

Centrohelid heliozoans, or Centroplasthelida Febvre-Chevalier et Febvre, 1984 are a monophyletic group of free-living, ubiquitous, predatory axopodial protists forming cysts (1). Centroplasthelida along with haptophytes constitute the Haptista supergroup (2). Centrohelid heliozoans are known for their remarkable diversity of external siliceous scales of different morphology (3–5). Centrohelids are characterized by worldwide distribution and inhabit freshwater (6, 7), marine (8), saline and hypersaline ecosystems (9, 10), as well as soil ecosystems (11). Centrohelids are known as cytotrophic protists that feed on small unicellular protists (flagellates, ciliates, algae) (3, 12), bacteria (13), and even cytotoxic cyanobacteria (14). Algivorous centrohelids are also known for their ability to establish associations with harbored algal cells (15, 16) and kleptoplasty (17, 18). However, despite over three centuries of microscopic observation, documented cases of centrohelid-bacterial associations remain scarce (17, 19, 20).

Over the extended evolutionary history, bacteria and algae have evolved a diverse array of strategies to survive the grazing pressure of protist predators (21–24). Ingested microorganisms can avoid lysosomal degradation in host cells (24) and establish associations that exhibit remarkable diversity, ranging from facultative to obligate and from mutualistic to parasitic (19, 25). Both endosymbiotic and ectosymbiotic relationships occur between protists, unicellular algae, and bacteria (25). Protist-bacterial associations are widely distributed across the tree of life, as protists from almost every taxonomic group form association with a wide range of prokaryotes (19, 24, 26, 27). Symbiotic relationships have been documented in most major supergroups and are particularly well described in ciliates (28–31) and amoebozoans (32–34). Mutualistic associations can confer benefits to the host via metabolite biosynthesis (35, 36) and energy production (37), thereby enabling protists to succeed or thrive in specific environments (35). Conversely, protists can function as “Trojan horses”, serving as environmental reservoirs that enhance bacterial persistence and maintain virulence traits (38). Furthermore, a growing body of research has highlighted the presence of human and animal pathogens within protist microbiomes (39–41).

Centrohelid heliozoans have rarely been the focus of studies on protist-bacterial interactions, with only two known reports to date (17, 20). Nevertheless, their potential role as microbial reservoirs has been indirectly highlighted through the presence of endosymbiotic *Chlorella* algae, which can host the *Acanthocystis turfacea* chlorella virus (ATCV-1) (42). Remarkably, DNA sequences of ATCV-1 have been detected in the human oropharyngeal virome and associated with cognitive changes in both humans and mice (43). Furthermore, despite *Chlorella heliozoae* being its only known host, ATCV-1 can persist in mouse macrophages and induce inflammatory responses (44). Thus, centrohelids may be involved in the circulation of biologically significant agents whose environmental role in health remains to be studied.

In the present study, we provide the first molecular data on the bacterial community composition of centrohelid heliozoans using high-throughput amplicon sequencing of the full-length bacterial 16S rRNA gene.

## MATERIAL AND METHODS

### Sampling site and collection

Samples containing centrohelid heliozoans were collected from aquatic biotopes. Three strains were isolated from planktonic water samples: two (G001 and G004) from oligotrophic Lake Teletskoe (West Siberia, Russia), and one (G065) from mesotrophic Lake Kuchak (West Siberia, Russia). Three strains (GB-2, G038, and G014) were isolated from water samples containing bottom sediment; and one (G007) from an artificial aquatic biotope in a fountain (Istanbul, Turkey). Planktonic water samples were collected using a 5-L plexiglass bathometer (Papanin Institute for Biology of Inland Waters Collection, RAS, Russia). Water samples containing bottom sediment were taken from the littoral zone after the direct rolling of the sediment. All samples were collected in sterile containers and stored at +4°C. For more detailed information on the sampling sites and established clonal cultures, see Table 1.

**TABLE 1.**
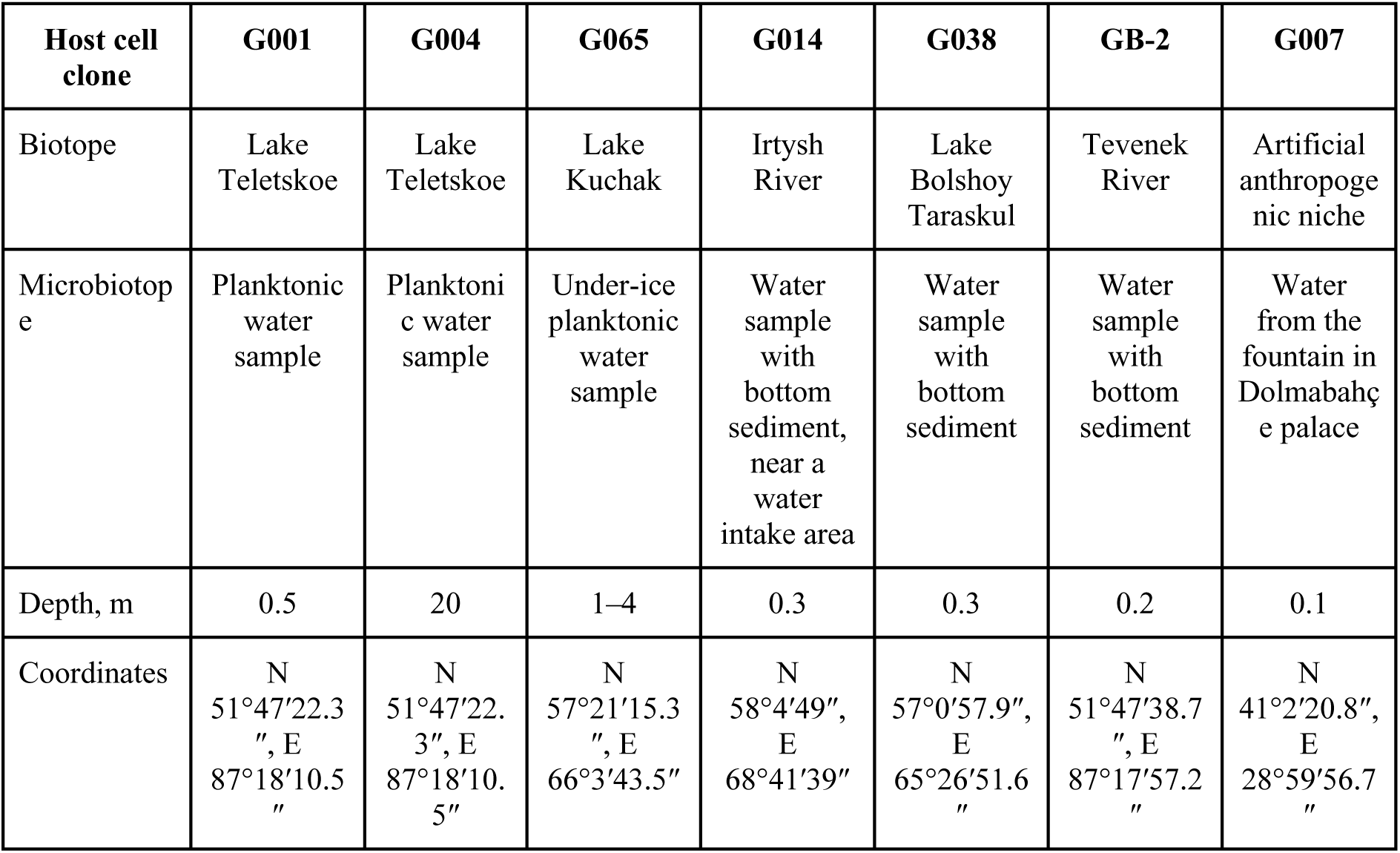
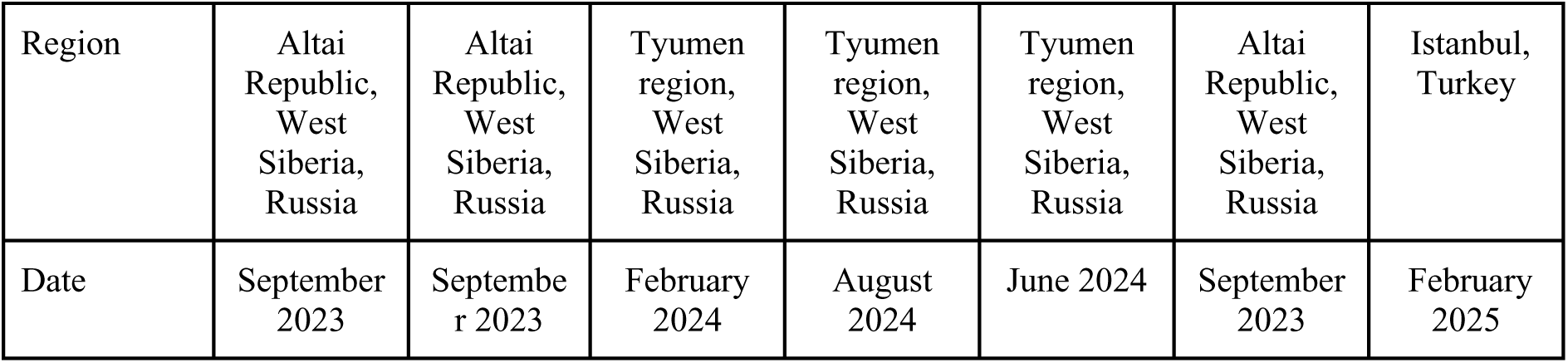
Characteristics of the sampling sites and clonal cultures.

**TABLE 2.** Morphological characteristics of the host cell cultures.

| Host cell clone | G001 | G004 | G007 | GB-2 | G014 | G038 | G065 |
| --- | --- | --- | --- | --- | --- | --- | --- |

| Taxa | *Acanthocystis* sp. 1 | *Acanthocystis* sp. 2/<br>*Heterophrys*-like organism | *Acanthocystis* sp. 3 | *Acanthocystis* sp. 4/<br>*Heterophrys*-like organism | *Raineriophrys* ssp. *sculpta* | *Heterophrys*-like organism | *Raineriophrys* sp./<br>*Heterophrys*-like organism |
| --- | --- | --- | --- | --- | --- | --- | --- |
| Diameter of living cells, $\mu\text{m}$ | 6.5–10.3<br>8.9±0.1<br>(n=52) | 8.4–26.5<br>14.9±0.7<br>(n=51) | 9.0–14.2<br>11.7±0.2<br>(n=55) | 14.8–23.1<br>18.7±0.7<br>(n=59) | 12.9–24.7<br>19.2±0.3<br>(n=96) | 9.8–16.9<br>12.7±0.2<br>(n=81) | – |

### Microscopy and host cell culture identification

For living cell observations and the establishment of heliozoan cell cultures, inverted Nexcope NIB950 and Nexcope NE950FL microscopes (Novel Optics, Ningbo, China) were used, equipped with differential interference contrast (DIC), phase contrast, and emboss contrast. An upright light microscope, the AxioScope A1 (Carl Zeiss, Jena, Germany), equipped with DIC and phase contrast oil objective (63×), was used to observe living cells and for photo and video documentation.

Centrohelids were identified based on morphological characteristics of their cell coverings using a Tescan Mira 3 scanning electron microscope (Tescan, Brno, Czech Republic). Preparation of the scales for scanning electron microscopy (SEM) was conducted according to Gerasimova and Plotnikov (45). For transmission electron microscopy (TEM), a suspension of live cells was transferred onto formvar-coated copper grids, air-dried, and washed with distilled water. The samples were observed using a JEM-1011 transmission electron microscope (JEOL, Tokyo, Japan) at an acceleration voltage of 80 kV.

### Clonal culture maintenance

The monoclonal cultures of the centrohelids from water samples were established by single-cell isolation and then maintained at +18°C in autoclaved, still Aqua Minerale water (PepsiCo, Inc., Moscow Oblast, Russia) inoculated with *Aeromonas sobria* bacteria and *Parabodo caudatus* flagellates strain BAS-1 (Papanin Institute for Biology of Inland Waters Collection, RAS).

Prior to microbiome study of heliozoan clonal cultures, we standardized the prey culture by isolating and cloning a single *P*. *caudatus* flagellate cell. A single cell of *P*. *caudatus* was repeatedly washed in sterile, still Aqua Minerale water to remove visible extracellular bacteria and was grown on Aqua Minerale water supplemented with *A. sobria* for 5 days at +18°C. Then, 2 mL of *P*. *caudatus* clonal culture was multiplied in 30 mL sterile Aqua Minerale water supplemented with *A. sobria* and cultivated for three days at +18°C. Subsequently, the 30 mL of *P*. *caudatus* clonal culture was aliquoted into three separate dishes, with 10 mL per dish, to constitute three biological replicates. To each of these three biological replicates, 3 to 5 centrohelid cells from the clonal culture, which had been repeatedly washed in a nuclease-free water (Qiagen, Germany), were added. The cultures were then incubated at +18°C for 10–14 days. Three biological replicates of *P. caudatus* flagellates cell cultures were used as a negative control for the centrohelid microbiome analysis.

### Metabarcoding library preparation and sequencing

For DNA analysis, 10 mL of the growth medium containing centrohelid, *P*. *caudatus* and *A. sobria* cells from each biological replicate was sequentially passed through 5.0 μm and 0.2 μm pore-size nitrocellulose membrane filters (Vladisart, Russia) using 25-mm syringe filter holders (Sartorius, Germany). Total DNA from each membrane filter was extracted by the ZymoBIOMICS DNA Miniprep Kit (Zymo Research, USA) according to the manufacturer’s protocol. DNA concentration was quantified with a Qubit 4 fluorometer (Invitrogen, USA) using a Qubit 1× dsDNA HS Assay Kit (Invitrogen, USA). DNA purity was assessed using an Implen NanoPhotometr (Implen, Germany). The full-length 16S rRNA gene was amplified using the primers 27F (5’–AGAGTTTGATCMTGGCTCAG–3’) and 1492R (5’–ACCTTGTTACGACTT–3’) (46) and sequenced on the Sequel II PacBio system following the SMRTbell library preparation protocol.

### Sequencing data analysis

Pacific Biosciences data were demultiplexed using the PacBio ‘lima’ program v. 2.5.1. HiFi reads (CCS reads with predicted accuracy ≥ Q20) were extracted using SAMtools v. 1.13 (47) and converted to FASTQ format using PacBio ‘bam2fasta’ v. 1.3.1. Further processing was performed with DADA2 v. 1.32.0 (48). Primer sequences were removed using the ‘removePrimers’ function, followed by quality filtering (maxEE=2), length selection (1000–1600 bp) and chimera removal. Error rates were learned independently for forward and reverse reads, and then the reads were merged. Taxonomy was assigned to the sequence variants using GTDB r220 database, up to the species level. Phyloseq v. 1.48.0 (49), ggplot2 (50), MicrobiotaProcess v. 1.16.0 (51), and microbiome v. 1.26.0 packages (52) were used for statistical analysis and visualization in R v. 4.4.0 environment. Alpha diversity indices (Chao1, Shannon, Gini–Simpson) were calculated using the microbiome R package. Beta diversity was assessed using Bray–Curtis dissimilarity. Principal coordinate analysis (PCoA) was performed, and the results were visualized using the ggordpoint function from the MicrobiotaProcess package. The statistical significance of group differences was tested with permutational multivariate analysis of variance (PERMANOVA).

Marker taxa were identified using linear discriminant analysis effect size (LEfSe) (53) as implemented in the microbiomeMarker package v.1.10.0 (54). The analysis was conducted on counts per million (CPM)-normalized amplicon sequence variant (ASV) abundance data. The significance threshold for the non-parametric factorial Kruskal–Wallis test among classes was set at *p* = 0.05. Subsequently, pairwise comparisons using the Wilcoxon test were performed, and p-values were adjusted using the false discovery rate (FDR) correction. Finally, the effect size was estimated using linear discriminant analysis (LDA), with an LDA score threshold of 2.0.

### Phylogenetic analysis of symbionts

Taxonomic classification and search for closest relatives were performed using the SINA aligner implemented in the SILVA ACT (Alignment, Classification and Tree) web service against the SILVA SSU Ref NR 99 database (release 138.2) (https://www.arb-silva.de/archive/release_138.2). To determine the precise evolutionary position of the target ASVs, three separate phylogenetic datasets were assembled. Each dataset included the analyzed ASVs, their closest reference neighbors retrieved from the SILVA database, and two selected outgroup sequences.

The Burkholderiaceae dataset comprised 16 nucleotide sequences with a final alignment length of 1511 bp. The Pseudomonadales dataset included 32 sequences (1517 bp alignment length), and the Rickettsiales dataset consisted of 23 sequences (1455 bp alignment length). Sequences were aligned using MAFFT ver. 7.475 with the L-INS-i algorithm (55).

Phylogenetic reconstructions were performed using Bayesian inference (BI) and Maximum Likelihood (ML) methods. Bayesian inference was performed using the parallel MPI version of MrBayes (56) under the GTR + I + G substitution model for all three datasets. Four independent runs with four Metropolis-coupled Markov chains each were run for 20 million generations, with a 50% burn-in. Convergence was assessed using MrBayes diagnostics, with the average standard deviation of split frequencies falling below 0.01. ML analyses were performed using IQ-TREE ver. 1.6.12 (57) with 1000 ultrafast bootstrap pseudoreplicates (58). The best-fit evolutionary models were determined using the built-in ModelFinder algorithm (59) and selected as TN+F+I+G4, TIM3+F+I+G4, and GTR+F+I+G4 for the Burkholderiaceae, Pseudomonadales, and Rickettsiales datasets, respectively.

## RESULTS

### Morphology of the host cells of centrohelid clones

The cells of all studied strains were spherical, with live cells measuring approximately 6.5–26.5 µm in diameter (Fig. 1). Axopodia were 2–4 times the length of the cell diameter. The cells of all the strains floated. Cells were observed floating freely in Petri dishes. Cyst formation was not observed during a year of maintenance in the live culture collection under laboratory conditions. The G065 strain clone died after 2 months of laboratory cultivation.

**FIG 1.**
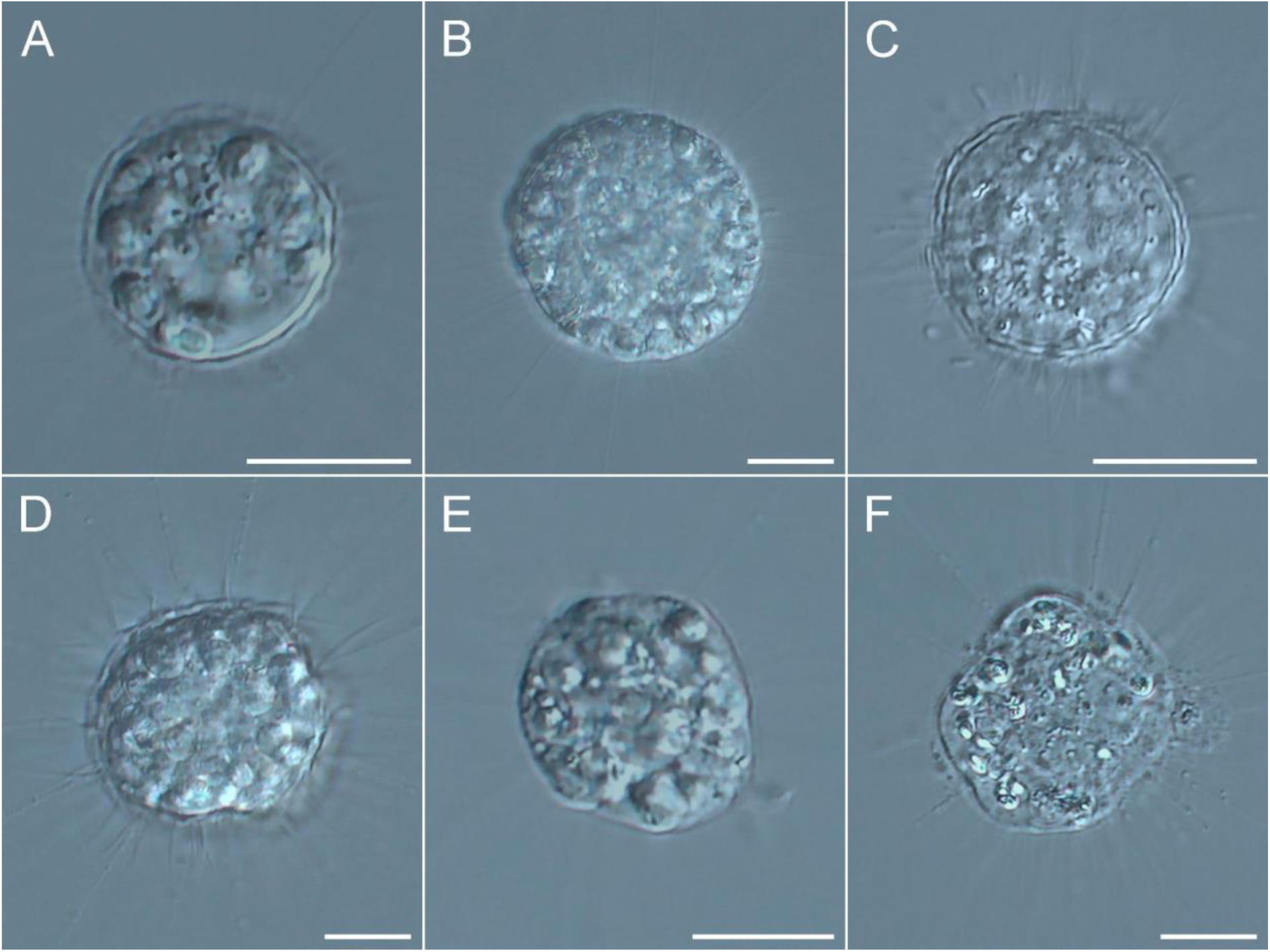
Morphology of living cells of investigated centrohelid clones, DIC (oil immersion), 100x. (A) Strain G001. (B) Strain G004. (C) Strain G007. (D) Strain G014. (E) Strain G038. (F) Strain GB-2. Scale bar: 5 μm.

Based on the morphology of the cell coverings (Fig. 2), we identified the taxonomy of the host cells.

**FIG 2.**
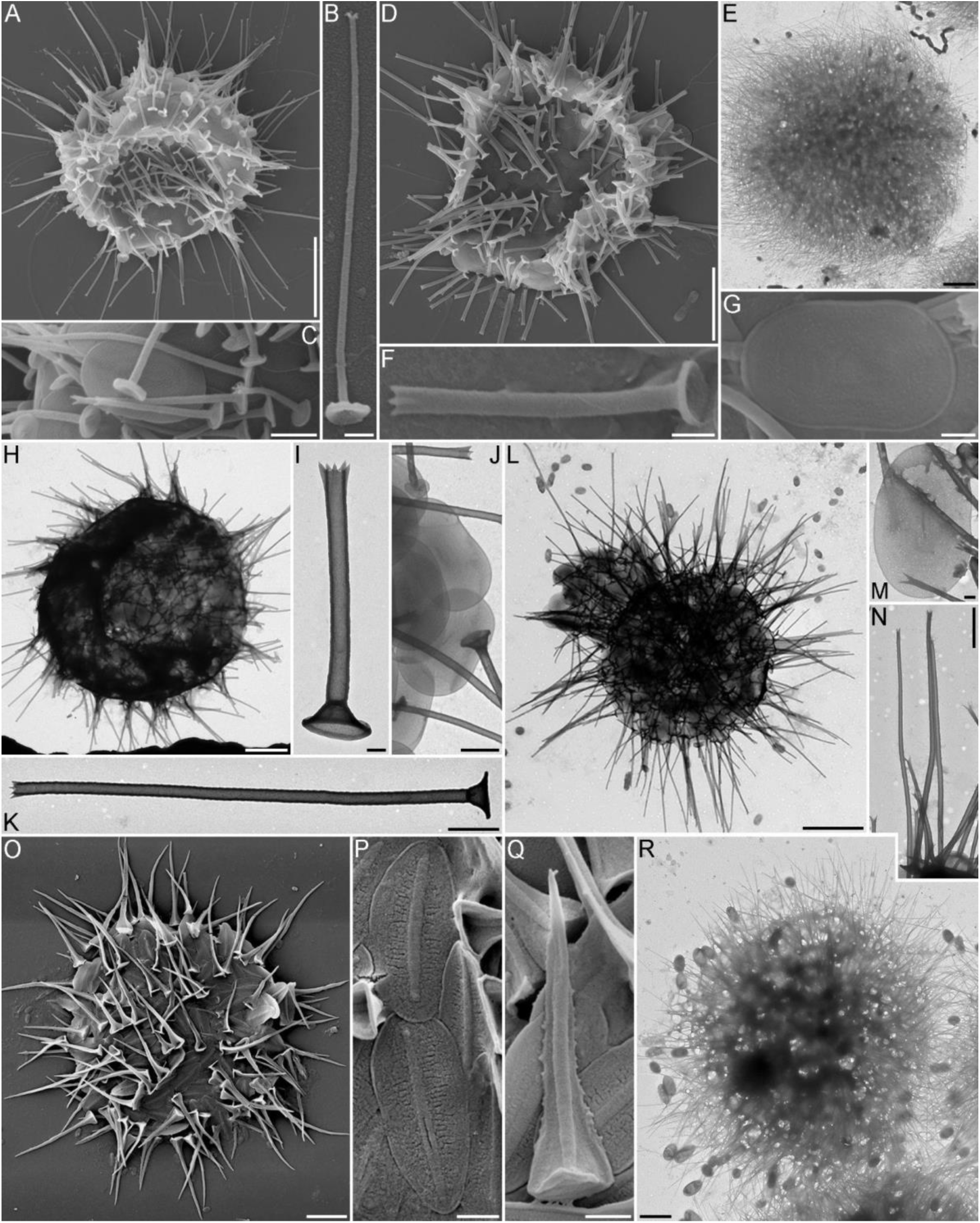
Morphology of the cell coverings of investigated centrohelid clones, SEM (A–G, O–Q), TEM (E, H–N, R). (A–C) *Acanthocystis* sp. 1, strain G001. (D, F, G) *Acanthocystis* sp. 2, strain G004. (E) *Heterophrys*-like stage of strain G004. (H–K) *Acanthocystis* sp. strain 4, GB-2. (L–N) *Acanthocystis* sp. 3, strain G007. (O–Q) *Raineriophrys* ssp. *sculpta*, strain G014. (R) *Heterophrys*-like organism, strain G038. (A, D, E, H, L, O, R) General view of dried cells. (B, F, I, K, N, Q) Spine-scales. (G, M, P) Plate-scales. (C, J) Scattered plate- and spine-scales. Scale bar: A, D, E, H, L – 5 μm; B, E, F, G **–** 0.5 μm; C, K, N, P, Q – 1 μm; I, M – 0.2 μm; J, R – 2 μm.

The cell coverings of strains G001 (Fig. 1A–C), G004 (Fig. 1D, F, G), GB-2 (Fig. 1H–K), G007 (Fig. 1L–N), and G014 (Fig. 1O–Q) were composed of both spine and plate scales. The plate scales of the strains G001, G004, GB-2, and G007 were oval or oviform (Fig. 1C, G, J, M). The spine scales consist of a shaft and circular basal plates ( (Fig. 1B, C, F, I, K). Shafts of the spine scales ended with teeth (Fig. 1B, F, I, K, N). Based on this morphology, the strains were assigned to the genus *Acanthocystis* Carter, 1863. The cells of strains G014 (Fig. 1O–Q) and G065 possessed plate and spine scales with pointed tips and triangular base plates and were assigned to the genus *Raphidocystis* Penard, 1904. However, strain G065 demonstrated a dimorphic life cycle, involving a change from spine scales (not visualized either in SEM or TEM images) to organic spicules, a characteristic of centrohelids grouped under the term *Heterophrys*-like organisms (HLO Bardele, 1975). Strain G038 was covered exclusively with spindle-shaped spicules, did not demonstrate a change in life cycle stages throughout the entire cultivation period, and was assigned to HLO (Fig. 1R).

### Sequence data overview

A total of 9,256,716 reads of the full-length 16S rRNA gene were obtained, of which 5,933,262 (64.1%) remained after quality filtering. The average depth of high-quality paired-end reads was about 128,983 per sample. Among the remaining 348 ASVs, 100% were assigned to phyla, 348 (100%) to classes, 348 (100%) to orders, 339 (97.4%) to families, and 336 (96.5%) to genus.

### Taxonomic composition of the microbial communities associated with the prey culture

Prior to analysing the centrohelid microbiome, we analysed the microbiome of the *P. caudatus* flagellate prey culture. The *P. caudatus* microbiome was predominantly composed of *Aeromonas sobria* (78% of total reads) and *Variovorax* sp. (21.8%), both assigning the phylum Pseudomonadota (Fig. 3). Other detected taxa, including *Sphingobium*, *Aureimonas*, and *Curvibacter* (all Pseudomonadota), as well as *Arcicella*, *Hymenobacter* (Bacteroidota), and *Prosthecobacter* (Verrucomicrobiota), were scarce, each constituting less than 0.1% of the total reads.

**FIG 3.**
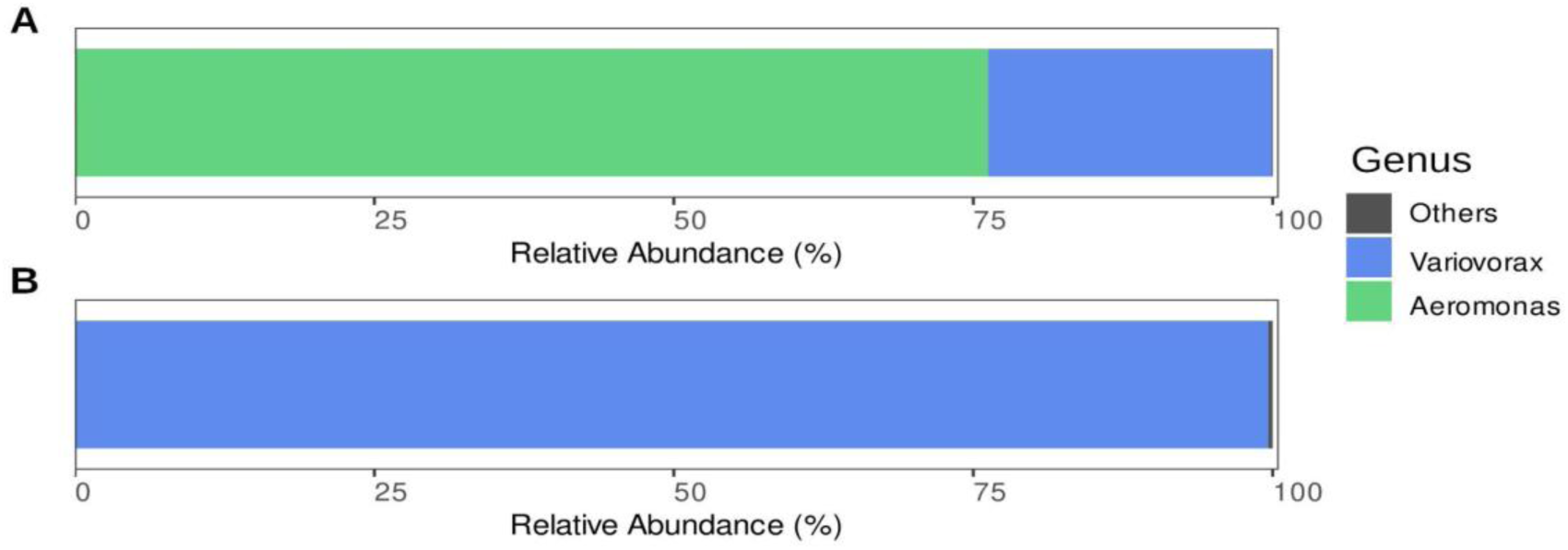
Taxonomic composition of bacterial communities associated with the *P. caudatus* flagellate prey culture at the genus level. (A) Relative abundances of dominant bacterial taxa (> 1%) in the prey culture fed *A. sobria* (corresponding to the genus name *Aeromonas*). (B) Relative abundances of dominant bacterial taxa (> 1%) of the prey culture after exclusion of *A. sobria* sequences.

The microbial community composition of the native *P. caudatus* microbiome was reanalysed following the exclusion of *A. sobria* sequences, which had been intentionally added as a food source. After removing these reads, no new major taxa appeared in the microbiomes, while the relative abundance of *Variovorax* sp. increased significantly to 99.8%. The relative abundances of the previously rare taxa (*Arcicella*, *Sphingobium*, *Aureimonas*, and *Prosthecobacter*) remained low.

Finally, all ASVs associated with the *P. caudatus* prey microbiome were excluded from the centrohelid microbiome analysis.

### Taxonomic composition of the microbial communities associated with the centrohelid strains

After excluding sequences from the prey culture microbiome, the centrohelid-associated microbiome comprised a total of 5 prokaryotic phyla, 6 classes, 22 orders, and 58 genera (Table S1). The community was dominated by the phyla Pseudomonadota (54.8% of total reads) and Bacteroidota (41.7%). The phylum Actinomycetota was scarce (3.5%), while Verrucomicrobiota, Bacillota, and other phyla were negligible, each representing less than 0.01% of the community (Fig. 4A).

**FIG 4.**
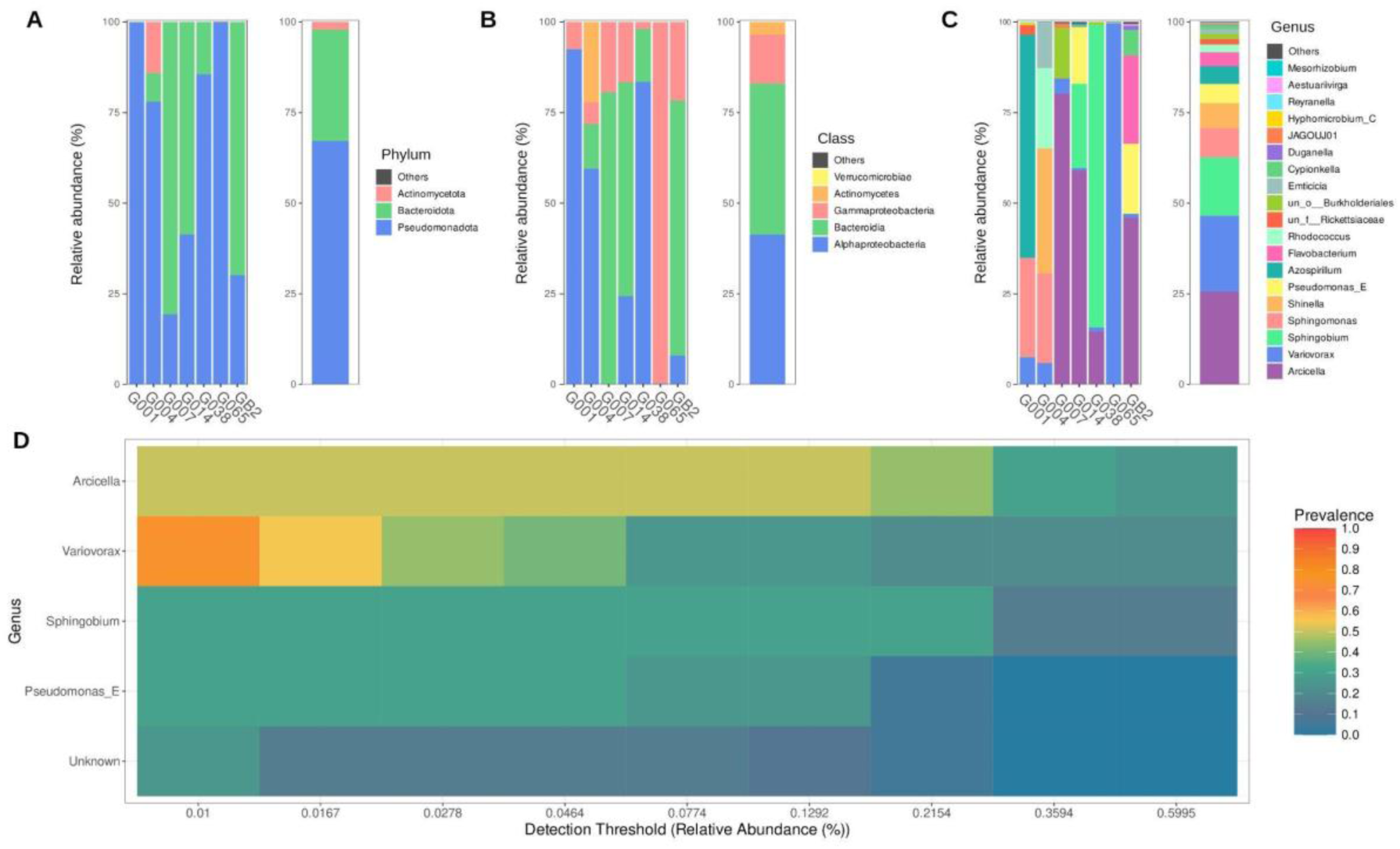
Taxonomic composition and core microbiome of bacterial communities associated with centrohelid cells. (A–C) Relative abundances of dominant bacterial taxa (> 1% in any sample) at the phylum (A), class (B), and genus (C) levels. The “Others” category represents the sum of all taxonomic groups with relative abundance below the 1% threshold. Color assignments correspond to specific taxon names. (D) Heatmap of the core microbiome at the genus level. Prevalence was assessed as ASV detected in the range from 50 to 100% samples.

The phylum Pseudomonadota dominated the microbiomes of strains G065 and G001, comprising almost 100% of the relative abundance in these two strains (Table S2). It also constituted 65% and 85% of the bacterial communities in strains G038 and G004, respectively (Fig. 4A). The microbiomes of strains G007, GB-2, and G014 were dominated by Bacteroidota (80.6%, 70.3%, and 59%, respectively), with Pseudomonadota representing 19–40% of the relative abundance. Actinomycetota was present only in the microbiome of strain G004, where it accounted for 22.2% of the relative abundance (Fig. 4A).

At the class level, Alphaproteobacteria and Bacteroidia were the most abundant, each accounting for 41% of the relative abundance, while Gammaproteobacteria represented 13.5% (Fig. 4B). Alphaproteobacteria were dominant in the microbiomes of strains G001 (92.5%), G038 (83.5%), and G004 (59.5%). In contrast, they accounted for 8% and 24% of the relative abundance in strains GB-2 and G014, respectively, and were absent from the G007 microbiome. Bacteroidia were absolutely dominant in the microbiomes of strains G007 (80.6%), G014 (59%) and GB-2 (70.3%). Members of this class also represented 12.4% and 14.6% of the community in strains G004 and G038 but were almost absent from strains G065 and G001. Gammaproteobacteria were overwhelmingly dominant in the microbiome of strain G065 (99.6%) and accounted for 16–21% in strains G014, G007, and GB-2 but less than 8% in strains G001, G004, and G038. Additionally, unassigned Gammaproteobacteria (order Burkholderiales) represented the second most abundant taxon in the G007 microbiome, accounting for nearly 14% of the relative abundance (Table S2). Phylogenetic reconstruction of 16S rDNA sequences (Fig. 5) grouped ASVs of the unassigned Burkholderiales (ASV265, ASV285, ASV292, ASV8, ASV200, ASV239, ASV243, ASV249, and ASV193) into a single clade with strong support (100% bootstrap, 1 posterior probability). The phylogenetic placement of this clade indicates a close relationship with the genus *Limnobacter*, suggesting that these sequences represent novel, previously undescribed taxa within this lineage.

**FIG 5.**
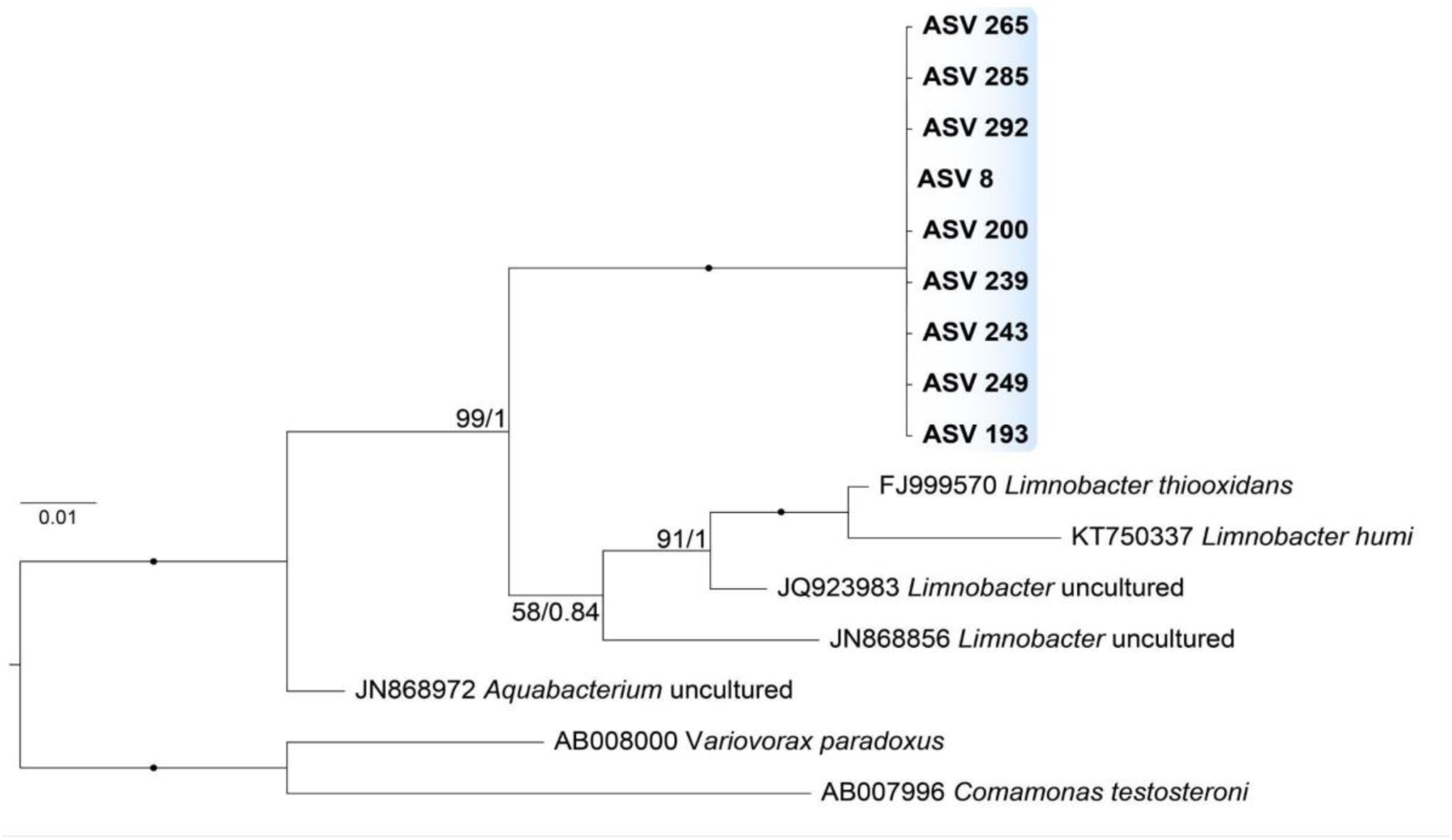
Maximum Likelihood phylogenetic tree showing the position of the recovered ASVs based on 16S rRNA gene sequences. Support values are indicated at nodes in the following format: Maximum Likelihood bootstrap percentage / Bayesian posterior probability. The tree is based on 1511 aligned nucleotide sites and includes 16 sequences. *Variovorax paradoxus* and *Comamonas testosteroni* were used to root the tree as an outgroup. Support values equal to 100/1.00 are indicated by solid black circles (●). The Maximum Likelihood analysis was performed using the TN+F+I+G4 substitution model, and Bayesian inference was conducted using the GTR+I+G model. Abbreviations: ASV, amplicon sequence variant.

At the genus level, *Arcicella* (34.7%) and *Sphingobium* (21.4%) were the dominant genera (Fig. 4C), followed by *Pseudomonas* (7.9%), *Sphingomonas* (6.5%), *Azospirillum* (5.8%), *Shinella* (5.5%), *Flavobacterium* (5.0%), *Variovorax* (3.7%), and *Rhodococcus* (3.5%). Several other genera, including *Emticicia* and *Cypionkella*, each accounted for approximately 2% of the relative abundance.

Notably, the genera *Arcicella*, *Variovorax*, *Sphingobium*, and *Pseudomonas* constituted the core microbiome (Fig. 4D). For instance, *Arcicella* was a core component in strains G007 (80.3% relative abundance), G014 (59%), and GB-2 (46%). *Variovorax* constituted the core microbiome of the G065 strain (99.6%). *Sphingobium* constituted the core microbiome of the G038 strain (83.5%). Finally, ASVs related to *Pseudomonas* were associated with the microbiomes of strains GB-2 and G014. Furthermore, despite the prevalence of *Arcicella*, *Variovorax*, and *Sphingobium* in the centrohelid microbiomes, the corresponding ASVs did not match those from the prey culture microbiome, even though these genera were present in both.

### Identification of opportunistic bacterial pathogens associated with centrohelid heliozoans

Among the 58 bacterial genera revealed in centrohelid microbiomes and their associated culture media, there are unexpectedly identified bacterial taxa recognized as opportunistic pathogens: *Acidovorax*, *Acinetobacter*, *Anaerococcus*, *Bosea*, *Corynebacterium*, *Escherichia*, *Moraxella*, *Mycobacterium*, *Prevotella*, *Pseudomonas*, *Ralstonia*, and *Sphingomonas*.

*Pseudomonas* and *Sphingomonas* were the most abundant genera, accounting for 7.9% and 6.5% of the relative abundance of total reads, respectively. Interestingly, *Pseudomonas* sp. constituted the core microbiome of the GB-2 strain accounting for 19.2% of relative abundance. Phylogenetic reconstruction of 16S rDNA sequences from pseudomonads revealed that individual ASVs formed well-supported clades with known pathogens (Fig. S1). ASV359, associated with the microbiomes of strains G001 and G014, formed a clade with three *P. aeruginosa* sequences (CP028162, CP030075, and ALBV02000037). ASV398, recovered from the microbiome of strain G065, formed a clade with sequences of *P. parafulva* CP019952 and *P. fulva* CP023048. Both clades were strongly supported (100% bootstrap support, 1 posterior probability).

*Sphingomonas* was mainly represented by ASV6, which remained unassigned at the species level and was associated with microbiomes of strains G001 and G004, accounting for 27.3% and 20.3%, respectively (Fig. 4C, Table S2). ASV21, identified as *Sphingomonas koreensis*, was associated with the microbiome of the strain G004, representing 4.3% of the relative abundance.

All other opportunistic pathogens (*Acidovorax temperans, Acinetobacter johnsonii*, *A*. *lwoffii*, *A*. *beijerinckii*, *Agrobacterium tumefaciens*, *Anaerococcus octavius*, *Corynebacterium sanguinis*, *Escherichia coli*, *Moraxella osloensis*, *Mycobacterium chlorophenolicum*, *Pseudomonas aeruginosa*, *Prevotella bivia*) were detected as minor components of the centrohelid microbiomes (Table S2). Nevertheless, the highest abundances of the aforementioned opportunistic pathogens were observed in strains G004 (*M. osloensis*, *A. lwoffii*, *R. pickettii*, and *C. sanguinis*) and G065 (*C. sanguinis*, *A. octavius*, *A. temperans*, and *E. coli*). All other centrohelids were associated with single taxa: strain G014 contained ASVs of *R. pickettii*, *E. coli*, and *P. aeruginosa*; strain G007 contained ASVs of *A. johnsonii* and *P. bivia*; strain GB-2 harbored ASVs of *E. coli* and *M. osloensis*; strain G001 was associated with ASV of *P. aeruginosa*.

Interestingly, members of Rickettsiaceae were also found in association with centrohelid microbiomes. ASV27, whose genus affiliation is uncertain, constituted 2.6% of the relative abundance in the microbiome of strain G001 (Table S2). ASV110 was identified as *Megaera polyxenophila* and was part of the microbiome of strain G014. Phylogenetic analysis based on 16S rDNA sequences confirmed that ASV27 and ASV110 were positioned within the family Rickettsiaceae (Fig. 6). The 16S rDNA sequence of ASV110 formed a clade with *Candidatus* Megaira polyxenophila AB688628. This clade was sister to *Candidatus* Megaira sequence FJ612282, and together they formed a well-supported clade (92% bootstrap support, 0.99 posterior probability) with other sequences of *Candidatus* Megaira (KT851814, JF429385, KC189769, JX105706, KT851825, EF667896, KT851816). The clade uniting all *Candidatus* Megaira sequences was strongly supported (100% bootstrap support, 1.0 posterior probability). ASV27 formed a clade with a Rickettsiaceae incertae sedis FPLK01001860 derived from metagenomic data and received strong Bayesian support (0.99 posterior probability) but moderate Maximum likelihood support (67% bootstrap).

**FIG 6.**
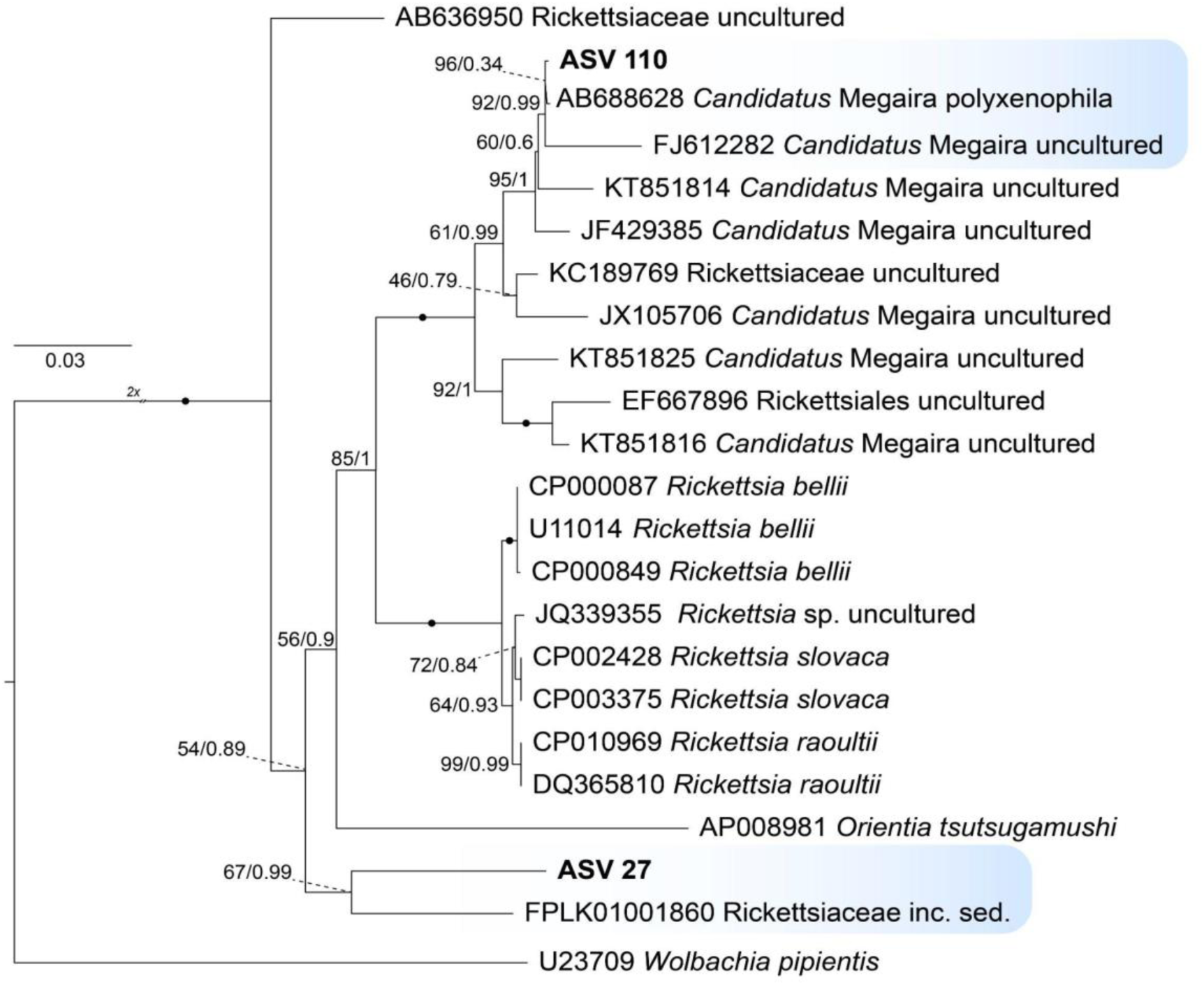
Maximum Likelihood phylogenetic tree showing the position of the recovered ASVs belonging to the family Rickettsiaceae based on 16S rRNA gene sequences. Support values are indicated at nodes in the following format: Maximum Likelihood bootstrap percentage / Bayesian posterior probability. The tree is based on 1455 aligned nucleotide sites and includes 23 sequences. *Wolbachia pipientis* was used to root the tree as an outgroup. Support values equal to 100/1.00 are indicated by solid black circles (●). The double slash (//) indicates that the branch length is reduced by half (2×). The Maximum Likelihood analysis was performed using the GTR+F+I+G4 substitution model, and Bayesian inference was conducted using the GTR+I+G model. Abbreviations: ASV, Amplicon Sequence Variant; inc. sed., incertae sedis.

### Comparative analysis of microbial communities between centrohelids cells and their culture media

To determine whether the microbiomes associated with centrohelid cells differed from those in the culture media, we used linear discriminant analysis effect size (LEfSe) method for a high-dimensional comparison. This analysis compared communities retrieved from centrohelid cells collected on 5-μm membranes with those from the culture medium collected on 0.2-μm membranes.

The LEfSe analysis revealed significant differences (p < 0.05) between the microbiomes of strains G014, G065, and GB-2 and their respective growth media. The relative abundance of the discriminating taxa changed by 4- to 5.5-fold (Table 3). For strain G014, the cell-associated microbiome was dominated by *Arcicella rosea* (Bacteroidia) and *Pseudomonas* sp. (Gammaproteobacteria), whereas its growth medium was predominantly composed of *Sphingobium* sp. and *Mesorhizobium* sp. (Alphaproteobacteria) (Table 3). In strain GB-2, the cell-associated microbiome was dominated by *Flavobacterium aquatile* (Bacteroidia) and members of the genus *Duganella* (Gammaproteobacteria). In contrast, its growth medium was primarily composed of *Arcicella rosea* (Bacteroidia), a notable difference from strain G014 along with *Variovorax* sp. (Gammaproteobacteria). For strain G065, the cell-associated microbiome was dominated by unclassified taxa within the phylum Pseudomonadota, which were 5.5-fold more abundant than in the medium.

**TABLE 3.** Bacterial taxa that differ significantly in abundance (LDA > 2, p < 0.05) between the microbial communities associated with centrohelid cells and their culture media.

| Host cell clone | Bacteria enriched in cells | Bacteria enriched in medium |
| --- | --- | --- |
| G001 | – | – |
| G004 | – | – |
| GB-2 | *Flavobacterium aquatile*<br>*Duganella* sp. | *Arcicella rosea*<br>*Variovorax* sp. |
| G007 | – | – |
| G014 | *Arcicella rosea*<br>*Pseudomonas*_E sp. | *Sphingobium* sp.<br>*Mesorhizobium* sp. |
| G038 | – | – |
| G065 | unclassified Pseudomonadota | *Variovorax* sp. |

In contrast, strains G001, G004, G007 and G038 exhibited no significant difference in bacterial diversity of the host cells and the culture medium.

### Microbiome diversity and specificity depending on host cell taxonomy and origin

To assess differences in microbiome complexity, we compared alpha and beta diversity metrics across centrohelid strains from different taxa and habitats (Fig. 7). Alpha diversity of the bacterial communities associated with heliozoan strains was consistently low (< 50 ASVs for most strains) and exhibited minimal variability among strains (Fig. S2). Differences between individual strains were not significant (p > 0.05). However, significant differences in Chao1 diversity were detected between *Acanthocystis* spp. (strains G001, G004, GB-2, G007) and the *Raineriophrys* spp. (G014, G065), as well as between *Heterophrys*-like organisms (G038) and the *Raineriophrys* spp. (G014, G065) (Fig. 7A). Habitat type did not significantly affect the alpha diversity of microbial communities associated with heliozoans (Fig. 7B). The only significant difference was observed between littoral and planktonic samples based on the Shannon index (p = 0.05).

**FIG 7.**
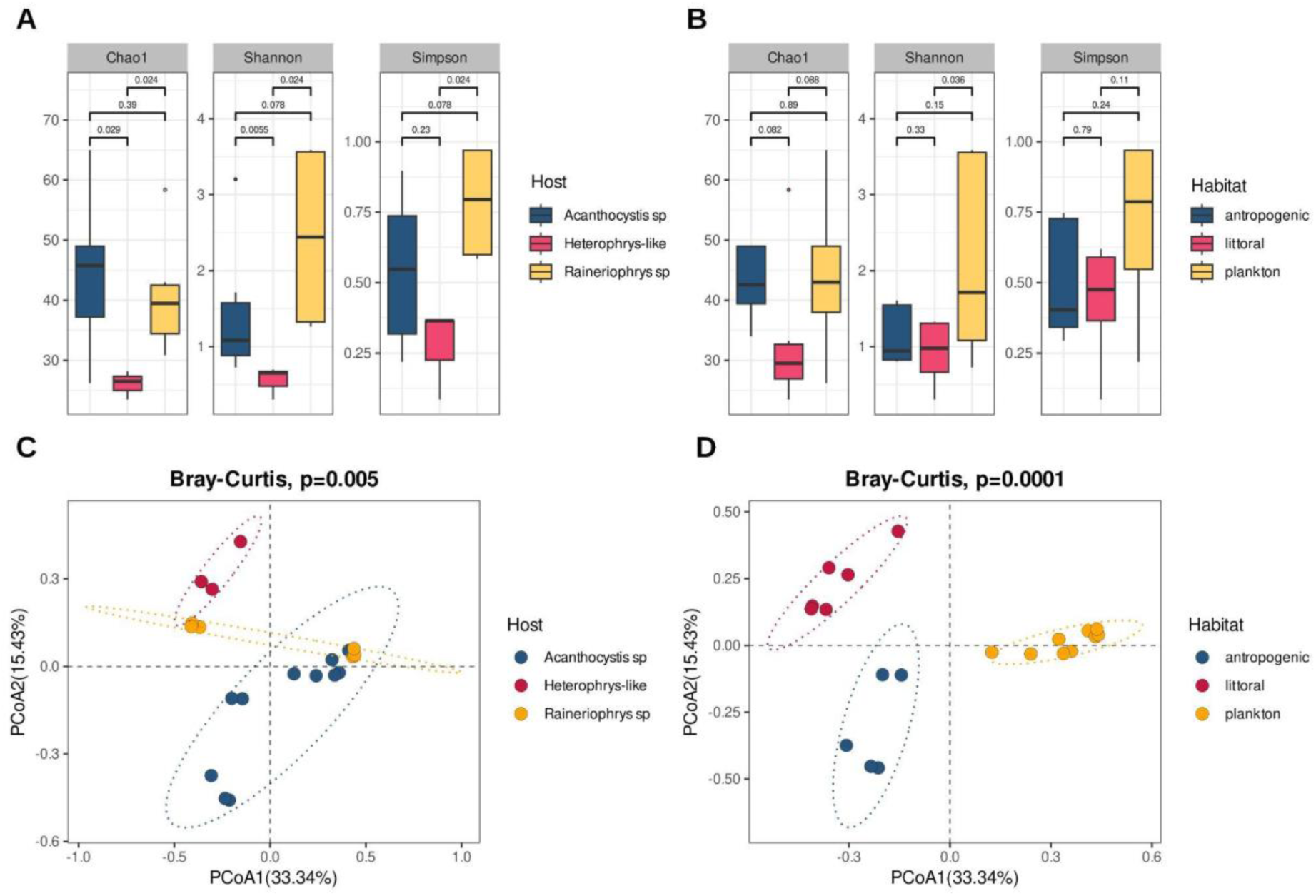
Alpha and beta diversity of bacterial communities associated with centrohelids. (A) Boxplot of alpha diversity indices (Chao1, Shannon and Simpson) for bacterial ASVs depending on host cell taxonomy. (B) Boxplot of alpha diversity indices (Chao1, Shannon and Simpson) for bacterial ASVs depending on habitat origin. (C) Beta diversity of bacterial communities associated with different centrohelid taxonomic groups. (D) Beta diversity of bacterial communities associated with centrohelids from different habitats. The ellipses represent the 95% confidence intervals for each group.

Beta diversity analysis, based on Bray–Curtis dissimilarity, revealed a significant effect of host species on microbial community composition (PERMANOVA: R² = 0.2361, F = 2.6271, *p* = 0.003). The members of the genus *Acanthocystis* (strains G001, G004, GB-2, G007) and the *Heterophrys*-like (strain G038) organisms formed distinct clusters, which merged with the cluster formed by the microbiome of the genus *Raineriophrys* (strains G014, G065) (Fig. 7C).

Habitat had an even stronger effect on community composition than host species (PERMANOVA: R² = 0.4457, F = 6.8357, *p* = 0.0001). The microbiomes formed three distinct clusters depending on their origin (Fig. 7D). Group 1 consisted of strains isolated from planktonic water samples of Lakes Teletskoe (Altai Republic, Russia) and Kuchak (Tyumen region, Russia) (G001, G004, and G065). Group 2 comprised strains G014 and G038, isolated from littoral zones containing water and bottom sediment. Group 3 included strain G007 from an artificial econiche (a fountain in Istanbul, Turkey) and strain GB-2 from a river water sample collected along an ecological trail (Altai Republic, Russia).

To further characterize differences in the microbial communities associated with centrohelid cells isolated from different habitats, we performed LEfSe analysis. This analysis identified 64 significant microbial biomarkers across the three habitat groups (Fig. 8). Among these, 6 taxa were characteristic of planktonic isolates, 14 of littoral zone isolates, while 55 taxa were significantly enriched in anthropogenic habitats. At the genus level, *Variovorax* was characteristic of planktonic isolates (G065, G001, G004). Several Alphaproteobacteria, namely *Sphingobium*, *Mesorhizobium*, CANJLN01 and *Megaera* were specific to littoral isolates (G014, G038). *Cypionkella*, *Aestuariivirga*, *Novosphingobium*, *Caulobacter*, *Asticcacaulis*, *Vitreimonas*, *Rhizobium*, UBA7672, and *Bosea* (Alphaproteobacteria), *Arcicella*, *Flavobacterium*, UKL13-3 (Bacteroidia), and *Duganella, Methylopumilus, Rhodoferax, Stutzerimonas,* JAUASW01 (Gammaproteobacteria) were specific for anthropogenic isolates (G007 and GB-2).

**FIG 8.**
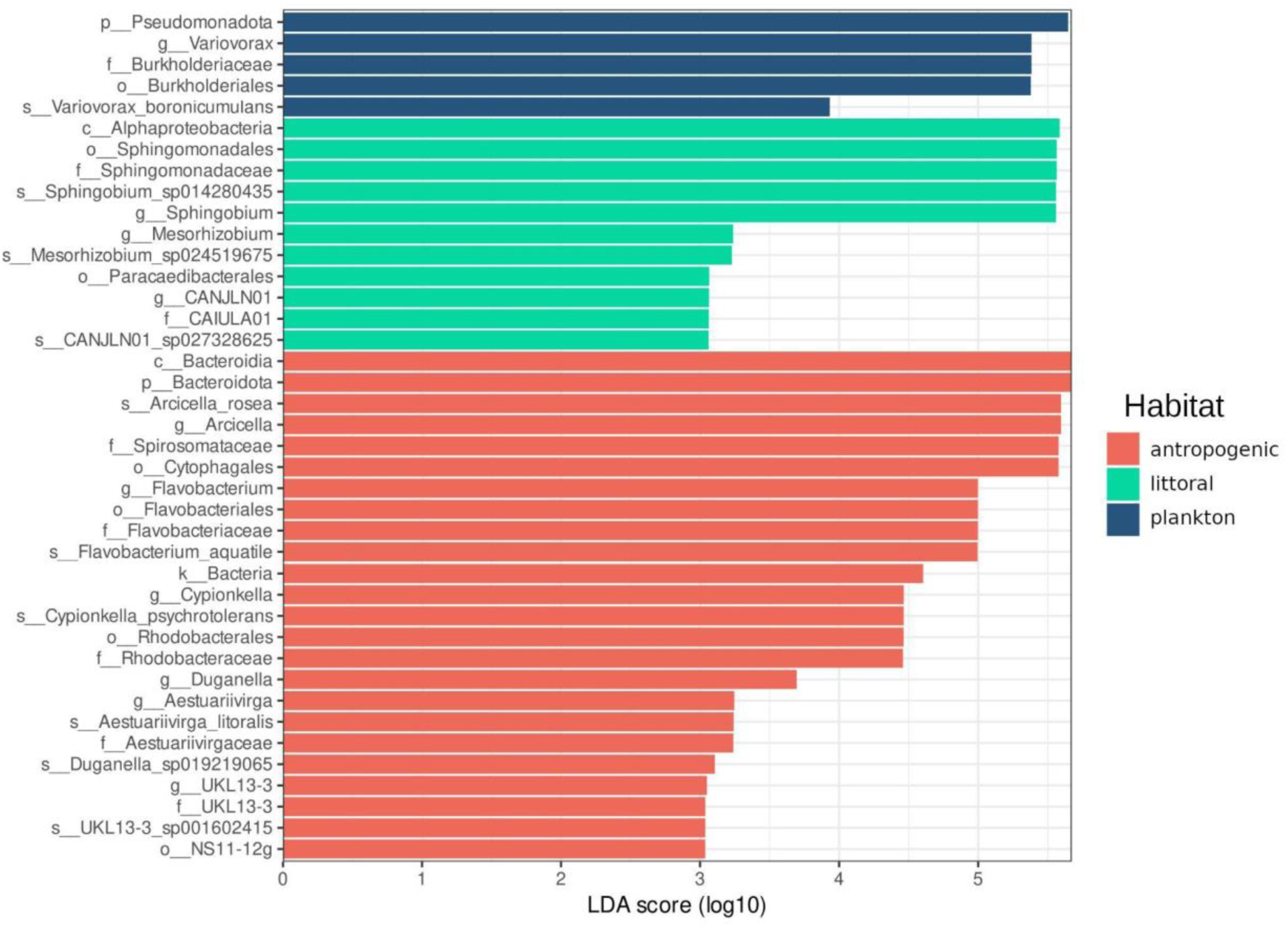
LEfSe analysis of microbial communities associated with centrohelid, isolated from anthropogenic, littoral and plankton habitats. LDA bar chart showing prokaryotic taxa with LDA score > 3.

## DISCUSSION

Protists are found in almost every environment on Earth (60), demonstrating remarkable adaptive capabilities that allow them to persist and flourish under drastically varied conditions (25, 61). Some of the most multifaceted adaptations involve symbiotic relationships with bacteria, archaea, unicellular algae, and other protists (19, 62, 63). Symbioses have been documented in most major supergroups and display different types of interactions (19, 25). While not ubiquitous, they remain well-studied in separate protist lineages, with particularly well-documented cases in amoebozoans (32–34) and ciliates (28–31). Recent reports from environmental isolates reveal novel associations, suggesting that symbiotic relationships are likely more widespread than traditionally assumed (64–66).

### Centrohelid microbiome composition

In this study, we performed, for the first time, a metabarcoding analysis using full-length 16S rDNA PacBio sequencing to investigate the microbial community associated with centrohelid heliozoans (Haptista, Centroplasthelida). We found that centrohelid strains from freshwater habitats were associated with bacteria from 5 phyla, 6 classes, and 58 genera. The metabarcoding analysis showed that Alphaproteobacteria, Bacteroidia, and Gammaproteobacteria were the most abundant classes in centrohelid microbiomes. Members of Alphaproteobacteria and Gammaproteobacteria are usually reported as the bacterial lineages most commonly found in symbiotic associations with free-living protists (19, 24, 67). In contrast, members of Bacteroidota have been mainly documented in host–bacterial associations with flagellated protists in termite guts, where they live in low-oxygen conditions and are involved in nitrogen fixation (68, 69). Bacteroidota have also been reported as dominant members of the microbiome in testate amoebae (70) and ciliates (71). Interestingly, in our study, members of the class Bacteroidia dominated the microbiome of centrohelids isolated from water samples with bottom sediments (G014, G038, GB-2), from a water sample taken from an artificial econiche (G007), and from a depth of 20 m (G004) (see Table S1 for more detailed information).

The genera *Arcicella*, *Sphingobium*, *Pseudomonas*, *Sphingomonas*, *Shinella*, *Azospirillum*, *Flavobacterium, Rhodococcus*, and *Variovorax* were identified as the most abundant bacteria within the centrohelid microbiomes. Among these, *Arcicella*, *Variovorax*, *Sphingobium*, and *Pseudomonas* constituted the core microbiome of centroheliozoa. Recently, *Pseudomonas*, *Variovorax*, *Azospirillum* and *Sphingomonas* have been reported as dominant genera in protist-associated bacterial communities in soil (72), in the microbiomes of free-living amoebae (40, 41, 73) and ciliates from water environments (74–76). In our study, *Pseudomonas* were mainly associated with the microbiomes of strains GB-2 (19.2%) and G014 (15.6%) and were scarcely represented in strains G001, G004, and G065. *Variovorax* ASVs overwhelmingly predominated the microbiome of strain G065 (99.6% relative abundance) and the prey culture of the flagellates *Parabodo caudatus* strain BAS-1 (99.8% relative abundance), however, they were represented by the different ASVs. Previously, *Variovorax* has been reported as a component of protist microbiomes (75) and was described as one of the most abundant taxa in the microbiome of the marine ciliate *Geleia* sp., accounting for up to 99% of the relative abundance among the most represented taxa (76). *Sphingomonas* have also been described as members of ciliate microbiomes (75) and were particularly revealed only in centrohelids from Lake Teletskoe (strains G001 and G004). Members of *Arcicella* are typically described as free-living inhabitants of aquatic ecosystems (77, 78), however, they have also been found to colonize fish skin (79) and the epidermis of ice worms (80). In our data, *Arcicella* were the dominant ASVs in the microbiomes of G007, G014, and GB-2, accounting for up to 79% of the relative abundance. The predominance of these genera across multiple strains suggests they may play important functional roles within the centrohelid microbiome.

### Prevalence of nitrogen-fixing bacteria in oligotrophic isolates suggests a potential role in nutrient acquisition

Some of the most striking patterns in community composition emerged from strains isolated from oligotrophic environments, where nutrient limitation may select for symbionts with specific metabolic capabilities. Interestingly, microbiomes of two clones (G001, G004) isolated from the oligotrophic Lake Teletskoe were dominated by the genera of nitrogen-fixing bacteria, such as *Azospirillum, Shinella,* and *Rhodococcus*. In strain G001, *Azospirillum* accounted for 56% of the relative abundance, while the microbiome of strain G004 was primarily composed of *Shinella* (34.4%) and *Rhodococcus* (22.2%). The genus *Azospirillum* is primarily known as a free-living, nitrogen-fixing, plant growth-promoting bacterium (81, 82). *Azospirillum* engages in complex interactions with other microorganisms. It promotes algal growth under salt stress (79) and, when co-inoculated with protists, enhances plant growth and nitrogen uptake (84). Recent research has expanded this understanding by revealing *Azospirillum* as a component of the microbiome of the free-living amoeba *Naegleria fowleri* (41). *Shinella* spp. are known symbionts of microalgae (85, 86) that can promote the growth and metabolism of their hosts. Furthermore, *Shinella* can convert nitrogen into ammonia, which may contribute to enhancing nutrient exchange between microalgae and bacteria (86). We also identified ASV of the denitrifying bacterium *Pelomonas puraquae* (ASV283) in the G004 strain microbiome, comprising almost 0.1% of the relative abundance. Previously, *P. puraquae* was isolated from sediments of oligotrophic waters and demonstrated significant aerobic denitrification capabilities (87). Additionally, *Pelomonas* sp. has been reported as a component of the free-living amoebae microbiomes (41). We also revealed other nitrogen-fixing bacteria, *Mesorhizobium* and *Azonexus*, in the microbiomes of strains G014 and G065, respectively, but they accounted for less than 0.1%. The prevalence of nitrogen-fixing bacteria in oligotrophic isolates raises the possibility that these symbionts may contribute to nutrient acquisition in nutrient-poor environments.

### Centrohelids as potential novel hosts for Rickettsiaceae

Symbiotic relationships are often functionally vital to the host, providing benefits through the biosynthesis of essential metabolites (19, 25) or by offering protection against pathogens (88). However, a growing body of evidence suggests that even symbionts engaged in evolutionarily established mutualistic relationships can act as parasitic or pathogenic symbionts, depending on the host they encounter and the prevailing environmental conditions (66). The family *Rickettsiaceae* (Alphaproteobacteria) represent a diverse group of obligate intracellular bacteria that infect a wide range of eukaryotic hosts, acting as symbionts, parasites, and pathogens (89, 90). In our study, members of *Rickettsiaceae* were detected in the microbiomes associated with two centrohelid strains. ASV110 was associated with the microbiome of strain G014 and, according to the sequence data analysis, was taxonomically related to *Candidatus* Megaira polyxenophila. In the phylogenetic tree of the family Rickettsiaceae (Fig. 6), ASV110 formed a well-supported clade with *Candidatus* Megaira polyxenophila AB688628, described as a symbiont of green algae *Carteria cerasiformis* (91), and was sister to the *Candidatus* Megaira sequence FJ612282 retrieved from lake sediment in China (92). The clade containing ASV110, AB688628, and FJ612282 grouped with other sequences of *Candidatus* Megaira, including those described as symbionts of the fish parasitic ciliate (KT851814) (93), an aquatic plant (KC189769) (94), the bacterioplankton community of the Dongjiang River in China (JF429385) (95), and a fish aquarium (JX105706). This clade was sister to a clade including *Candidatus* Megaira sequences isolated from parasitic ciliate (KT851825, KT851816) (93) and the epithelium of *Hydra oligactis* (EF667896) (97). *Candidatus* Megaira is an interesting bacterial symbiont that is phylogenetically closely related to the pathogen *Rickettsia* (39). Recently, *Candidatus* Megaira has been found in ciliates, amoebae, chlorophyte and streptophyte algae, as well as cnidarians (39). We additionally revealed an ASV27 belonging to an unassigned genus within the family Rickettsiaceae in association with strain G001. In the phylogenetic tree, ASV27 formed a cluster with Rickettsiaceae incertae sedis FPLK01001860, a sequence derived from metagenomic data originating from a freshwater sample (ENA project PRJEB17706). This clade was sister to *Orientia tsutsugamushi* AP008981 and a clade uniting sequences of *Rickettsia bellii* (CP000087, U11014, CP000849), uncultured *Rickettsia* sp. (JQ339355), *R*. *slovaca* (CP002428, CP003375) and *R*. *raoultii* (CP010969, DQ36581). These findings significantly expand the known host range for Rickettsiaceae and suggest that centrohelids may serve as previously unrecognized hosts for these parasitic bacteria.

### Centrohelids as potential reservoirs for pathogens

The presence of Rickettsiaceae raises broader questions about the potential role of centrohelids in harboring bacteria with clinical relevance. Unexpectedly, in the centrohelid microbiomes, we detected bacterial taxa belonging to opportunistic pathogens, including *Acidovorax*, *Acinetobacter*, *Anaerococcus*, *Bosea*, *Corynebacterium*, *Escherichia*, *Moraxella*, *Mycobacterium*, *Prevotella*, *Pseudomonas*, *Ralstonia*, and *Sphingomonas*. Within these genera, individual ASVs were identified to the species level and were related to *Sphingomonas koreensis*, *Moraxella osloensis*, *Acinetobacter lwoffii*, *A*. *johnsonii*, *A*. *beijerinckii*, *Pseudomonas aeruginosa*, *Ralstonia pickettii*, *Corynebacterium sanguinis*, *Anaerococcus octavius*, *Acidovorax temperans*, *Mycobacterium chlorophenolicum*, *Prevotella bivia*, and *Escherichia coli*. Among them, *S. koreensis* (ASV21) was the most abundant ASV, revealed in the microbiome of clone G004, isolated from oligotrophic Lake Teletskoe at 20 m depth. *S*. *koreensis* was first isolated from natural mineral water in Korea (98) and has since been rarely reported as a human pathogen. However, few cases have described *S. koreensis* as a causative agent of meningitis (99) and as an opportunistic human pathogen in hospitalized patients (100). Overall, the highest abundance of opportunistic pathogenic bacteria was observed in the microbiomes of clones from planktonic water samples, specifically G004 (*M. osloensis*, *A. lwoffii*, *R*. *pickettii*, and *C. sanguinis*) and G065 (*C. sanguinis*, *A. octavius*, *A. temperans* and *E. coli*). ASVs of *R*. *pickettii* and *E. coli* were also detected in the microbiome of G014 heliozoans. The microbiome of the clone G007 contained ASVs of *A. johnsonii* and *P. bivia,* and clone GB-2 included ASVs of *E. coli* and *M. osloensis*. ASVs of *P*. *aeruginosa* were detected in insignificant amounts in the microbiomes of strains G001 and G014. Recently, *R. pickettii*, *E. coli* and *P. aeruginosa* have been shown to be extracellular members of free-living amoebae with potential or established pathogenicity (101–103). Low-abundance ASVs corresponding to the genus *Bosea* were detected in the microbiomes of strains G004 and GB-2. Members of this genus are known for their resistance to digestion by free-living amoebae (21).

Free-living amoebae and ciliates are known to serve as an important ecological niche for pathogenic bacteria (73, 104, 105), protecting them from environmental stressors and increasing their transmission (106). Acting as “Trojan horses” of the microbial world (38), they also function as evolutionary “training grounds” where bacteria acquire enhanced virulence and antibiotic resistance after passage through protists (105, 107–109).

The majority of opportunistic pathogens detected in centrohelid microbiomes have been previously isolated from various environmental sources (110), as well as from hospital settings and clinical specimens (111–113). While some of these taxa are considered commensals with minor clinical significance (114) or have not previously been associated with human infections (115), others have been recovered from clinical specimens and linked to a range of human diseases. These include nosocomial infections (114, 116, 117), bacteremia (115, 118, 119), and peritonitis (120, 121). Moreover, *A. temperans* have even been suggested to play a functional role in lung cancer development (122). Although pathogens were found to be minor components of the centrohelid microbiome, this finding should not be neglected, as even a few cells of a virulent strain may be sufficient to cause disease. While ciliates and free-living amoebae are recognized as natural reservoirs for human pathogens (39–41, 74, 123), this study provides the first evidence that heliozoans may also harbor specific pathogens, expanding the known repertoire of protist hosts for clinically relevant bacteria.

### Factors shaping alpha and beta diversity of the bacterial community, associated with centrohelids

Modern research convincingly demonstrates that the structure of microbial communities is determined by a complex interplay of various factors, with interspecific interactions playing a primary role. Specifically, an analysis of microorganisms in European freshwater lakes has shown that biotic factors such as trophic interactions between bacteria and protists contribute significantly to the formation of community structure, surpassing the importance of physico-chemical environmental variables (124). Studies on rhizosphere communities in the Tibetan Plateau further confirm this, revealing that bacteria serve as central links (“bridges”) between different microbial groups (bacteria, protists and fungi), and the presence of predatory protists enhances overall system stability (125). However, resistance to phagocytosis is not the primary factor shaping the bacterial microbiota of protists (126, 127). Instead, environmental variability and host specificity have been identified as key drivers of bacterial community composition and its temporal variation in protist microbiomes (126, 128).

In our study, we detected that the bacterial community associated with centrohelids was shaped by host taxonomy (PERMANOVA, *p* = 0.005) and, even more strongly, by habitat origin (PERMANOVA, *p* = 0.0001). Indeed, *Sphingobium* was more abundant in *Heterophrys*-like strains, while *Mesorhizobium*, *Megaera*, and CANJLN01 were associated with *Raineriophrys* spp. strains. However, a LEfSe analysis revealed a significantly larger number of taxa (64) were specific to habitat origin. Both factors were thus identified as key determinants shaping the microbial composition associated with centrohelids in our study.

Overall, alpha diversity (richness) was relatively low (mostly <50 ASVs per strain) and showed little variation between different host strains. A similar pattern was observed in social amoebae *Dictyostelium discoideum* isolated from soil, where lower alpha diversity in the microbiome compared to the surrounding environment was attributed to host selectivity and the small size and restricted habitat provided by the amoebae (126). Furthermore, the lack of variation across different habitats in our study indicates that environmental factors do not significantly influence the richness of the associated bacterial communities. Taken together, the formation of protists microbiome is a multifactorial process, in which trophic interactions, complex networks of interspecific relationships, as well as physico-chemical conditions, and geographical location play a key role.

## Conclusion

This study provides the first comprehensive insight into the bacterial microbiota of centrohelid heliozoans and demonstrates that both host taxonomy and habitat origin contribute to community assembly. The detection of taxa with known pathogenic potential, together with the previously documented role of centrohelids as viral carriers, raises questions about their possible involvement in the environmental persistence and dispersal of microbes relevant to human health. Future research should explore the functional nature of these associations and assess whether centrohelids can serve as vectors or reservoirs for opportunistic pathogens in natural and anthropogenic ecosystems.

## Acknowledgments

The authors are grateful to Denis Tikhonenkov for providing the prey culture of the flagellates *Parabodo caudatus* BAS-1 strain, maintained in the living culture collection of the IBIW RAS. E.A.G. thanks Mikhail Soloviev and Grigory Romanenko for their invaluable help with the sampling on Lake Teletskoe.

## AUTHOR CONTRIBUTIONS

Elena A. Gerasimova: Conceptualization, Methodology, Investigation, Resources, Funding acquisition, Writing – original draft. Alexander S. Balkin: Investigation, Data curation, Validation, Visualization, Formal analysis, Writing – review & editing. German A. Sozonov: Data curation, Methodology, Investigation, Formal analysis, Visualization. Tatyana A. Chagan: Investigation, Formal analysis. Elizaveta I. Kaleeva: Investigation, Formal analysis. Ruslan Kasseinov: Investigation, Visualization. Darya V. Poshvina: Methodology, Investigation, Formal analysis, Writing – review & editing.

## Data availability

All raw sequencing reads for metabarcoding data have been deposited in the Sequence Read Archive (SRA) under project number (#to be released prior to publication). The prokaryotic 16S rRNA sequences generated in this study have been deposited in NCBI GenBank under accession numbers (#to be released prior to publication).

## Supplemental Material

**FIG S1.**
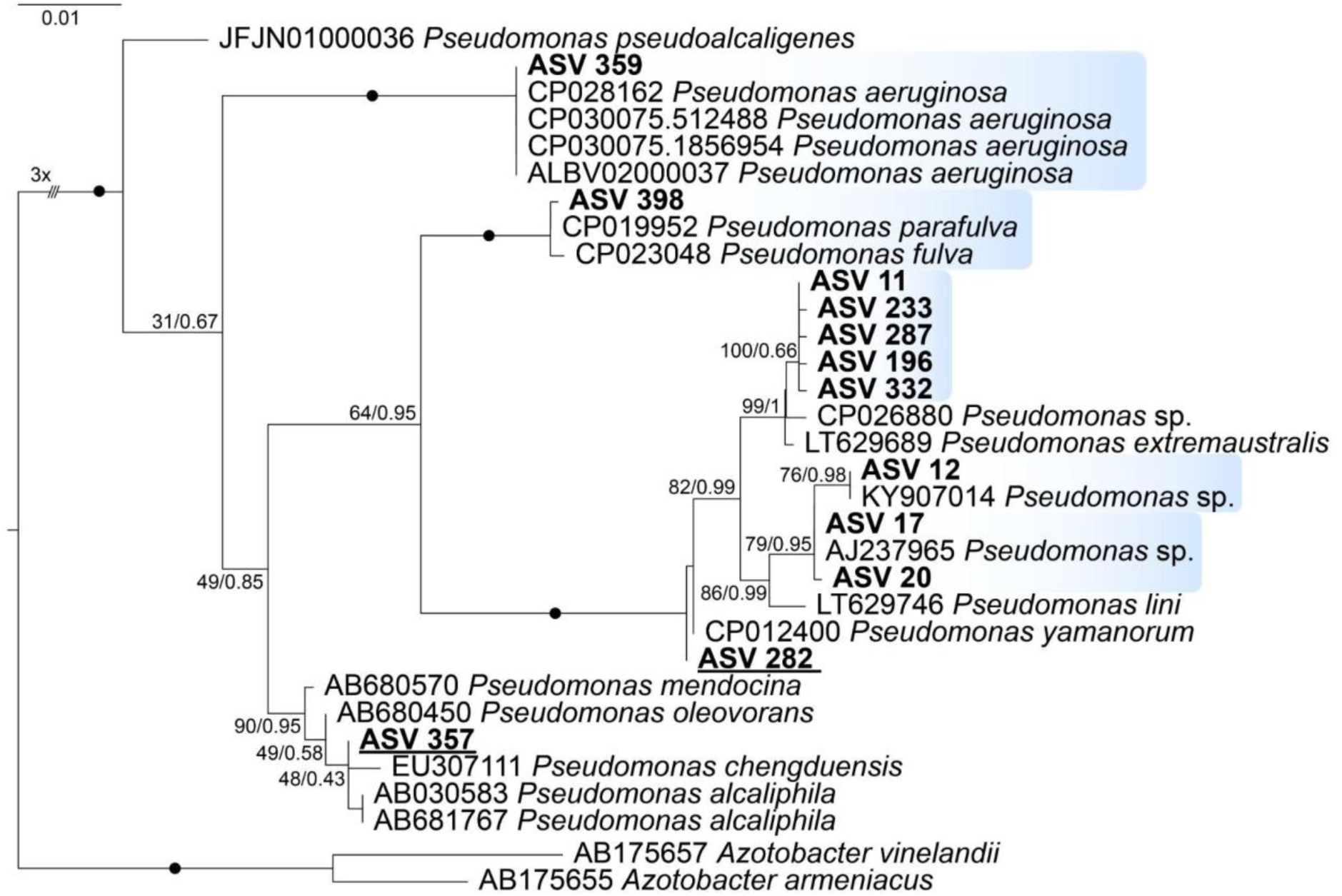
Maximum Likelihood phylogenetic tree showing the position of the recovered ASVs belonging to the genus *Pseudomonas* based on 16S rRNA gene sequences. Support values are indicated at nodes in the following format: Maximum Likelihood bootstrap percentage / Bayesian posterior probability. The tree is based on 1517 aligned nucleotide sites and includes 32 sequences. *Azotobacter vinelandii* and *Azotobacter armeniacus* were used to root the tree as an outgroup. Support values equal to 100/1.00 are indicated by solid black circles (●). The triple slash (///) indicates that the branch length is reduced by a factor of three (3×). The Maximum Likelihood analysis was performed using the TIM3+F+I+G4 substitution model, and Bayesian inference was conducted using the GTR+I+G model. Abbreviations: ASV, Amplicon Sequence Variant.

**FIG S2.**
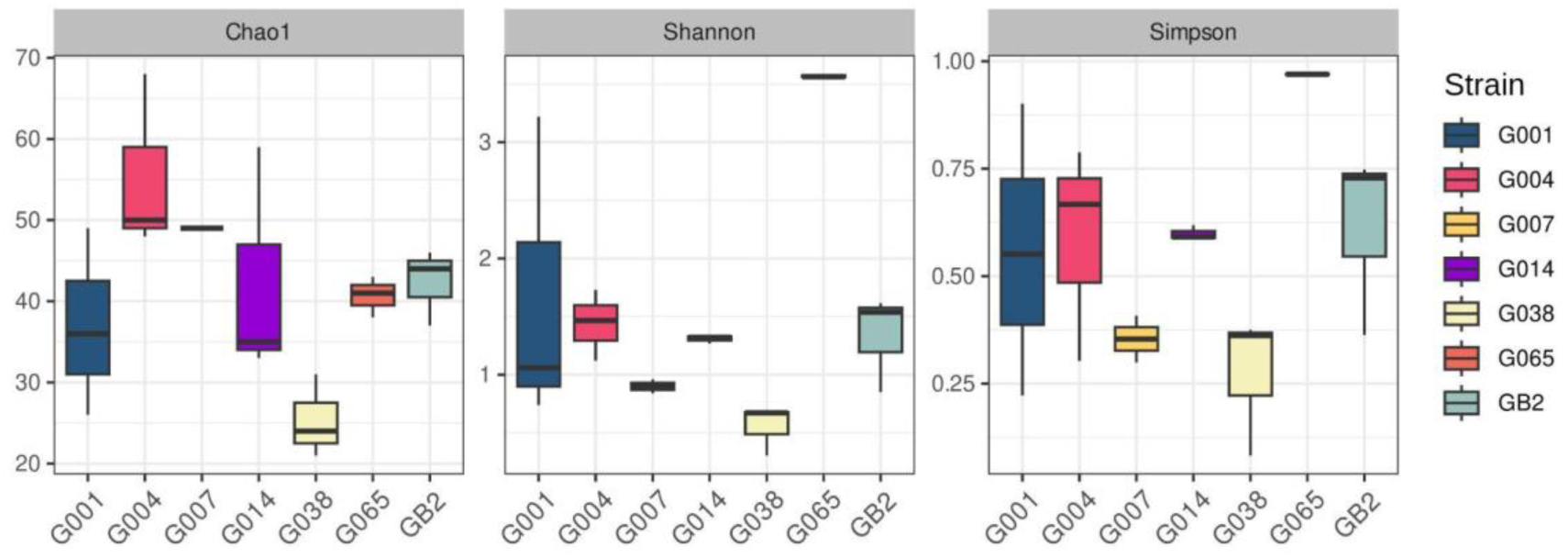
Alpha diversity of bacterial communities associated with centrohelids. Boxplots show the distribution of Chao1, Shannon, and Simpson indices for each individual strain.

## Abbreviations

ASV: amplicon sequence variant
DIC: differential interference contrast
LDA: linear discriminant analysis
LEfSe: linear discriminant analysis effect size
ML: Maximum likelihood
PCoA: principal coordinate analysis
SEM: scanning electron microscopy
TEM: transmission electron microscopy

